# Light Sheet Fluorescence Microscopy as a New Method for Unbiased Three-Dimensional Analysis of Vascular Injury

**DOI:** 10.1101/2020.01.02.893065

**Authors:** Nicholas E. Buglak, Jennifer Lucitti, Pablo Ariel, Sophie Maiocchi, Francis J. Miller, Edward S. M. Bahnson

## Abstract

**Aims:** Assessment of preclinical models of vascular disease are paramount in the successful translation of novel treatments. The results of these models have traditionally relied on 2-D histological methodologies. Light sheet fluorescence microscopy (LSFM) is an imaging platform that allows for 3-D visualization of whole organs and tissues. In this study, we describe an improved methodological approach utilizing LSFM for imaging of preclinical vascular injury models while minimizing analysis bias.

**Methods and Results:** The rat carotid artery segmental pressure-controlled balloon injury and mouse carotid artery ligation injury were performed. Arteries were harvested and processed for LSFM imaging and 3-D analysis, as well as for 2-D area histological analysis. Artery processing for LSFM imaging did not induce vessel shrinkage or expansion, and was reversible by rehydrating the artery, allowing for subsequent sectioning and histological staining *a posteriori*. By generating a volumetric visualization along the length of the arteries, LSFM imaging provided different analysis modalities including volumetric, area, and radial parameters. Thus, LSFM-imaged arteries provided more precise measurements compared to classic histological analysis. Furthermore, LSFM provided additional information as compared to 2-D analysis in demonstrating remodeling of the arterial media in regions of hyperplasia and periadventitial neovascularization around the ligated mouse artery.

**Conclusions:** LSFM provides a novel and robust 3-D imaging platform for visualizing and quantifying arterial injury in preclinical models. When compared with classic histology, LSFM outperformed traditional methods in precision and quantitative capabilities. LSFM allows for more comprehensive quantitation as compared to traditional histological methodologies, while minimizing user bias associated with area analysis of alternating, 2-D histological artery cross-sections.

**Translational Perspective:** A more reproducible and robust quantitation of vascular pathology in preclinical models is necessary to accelerate translational discovery. Current methodology to assess vascular disease has significant limitations. The methodology described herein employs a modern imaging modality, light sheet fluorescence microscopy (LSFM), to improve assessment of established preclinical vascular injury models. LSFM provides more comprehensive and precise analysis capabilities than classical histological approaches. Hence, LSFM applied to vascular research has the potential to drive new basic discoveries, and ultimately translation of novel therapies.

## I. Introduction

Cardiovascular disease (CVD) is the leading cause of death and disability in the world^1, 2^. Atherosclerosis is the major underlying cause of most CVD. Severe, symptomatic atherosclerosis is treated by percutaneous or surgical revascularization, and the long-term success of both approaches is limited by arterial restenosis^3^. Despite the advances in revascularization procedures, restenosis rates remain unacceptably high^3^. Hence, animal models are essential to understand the pathophysiology of restenosis and to discover and optimize new drugs to improve surgical outcomes. Restenosis occurs secondary to the development of neointimal hyperplasia and vessel remodeling^4^. Several preclinical injury models exist for studying restenosis^5^ including the rat carotid artery balloon injury model and the mouse carotid ligation model.

Classically, histological methods are used to quantify neointimal hyperplasia in preclinical models^6^. This classic approach involves sectioning the artery into two-dimensional (2-D) cross-sections and staining a selection of sections for analysis. A major limitation with this approach is that it assumes predictable and homogenous arterial injury for each animal and between animals. This suggests that a subset of microns-thick cross-sections will represent the lesion as a whole. However, arterial injury is not homogenous^7^, which is also observable using the methodology described herein. Furthermore, the pathology of neointimal hyperplasia is defined, but no consensus exists on how to properly quantify it. The literature varies between using three to ten cross-sections at variable regions throughout the artery for quantification^8-23^, which has the potential to introduce bias based on which and how many slices are analyzed. Moreover, classic histology limits our ability to observe and measure vessel remodeling, the other major component of restenosis, because only a few cross-sections are used to inform all of the morphological changes in the artery.

Light sheet fluorescence microscopy (LSFM) allows fast imaging of large (several mms in diameter), fixed, transparent tissue, by illuminating samples with a sheet of laser light from the side, and collecting the resulting fluorescence from the excited plane at a 90° angle^24^. This technique generates optical sections of the sample, is significantly faster, and causes less photo damage compared with other commonly used optically sectioning techniques such as laser scanning confocal and multiphoton microscopy^25^. This powerful tool has been used in a wide range of applications from imaging intracellular dynamics in a single cell, to developmental dynamics in entire organisms^26^. Indeed, LSFM has unraveled new mechanisms for megakaryocyte function and platelet migration from the bone marrow^27, 28^. Herein we describe a methodology using LSFM to image preclinical restenosis murine models in three-dimensions (3-D). We quantify changes in neointimal hyperplasia and arterial media volumes, alongside vessel remodeling, in response to arterial ligation or endothelial denudation. Additionally, we benchmark this novel methodology against classic histological analysis. To image the arteries, they must be made transparent by homogenizing their refractive index. Out of the many clearing methods available^29^, we selected iDISCO+^30^ for its excellent clearing, speed, low cost, and—critically— optimization for immunostaining in large tissue samples. We believe utilizing LSFM for analysis of preclinical restenosis models will improve rigor and reproducibility associated with these models and may identify novel mechanisms of this pathology. The overall aim of this manuscript is to provide a framework for analyzing preclinical restenosis models.

## II. Methods

All rat procedures described here have been approved by the Institutional Animal Care and Use Committee (IACUC) of the University of North Carolina at Chapel Hill. All mouse procedures were approved by the Durham Veterans Affairs Medical Center (DVAMC) IACUC (2013-002). All procedures conform to the Guide for the Care and Use of Laboratory Animals published by the United States National Institutes of Health.

### Reagents and Materials

Acetic Acid (ARK2183-1L, Sigma-Aldrich St. Louis, MO). Dibenzyl ether (DBE) (33630-1L, Sigma-Aldrich). Dichloromethane (DCM) (BDH1113-4LG, VWR, Radnor, PA). EDTA (BDH9232, VWR). Methanol (A4525K, Thermo-Fisher Scientific, Waltham, MA). Microcentrifuge tubes (05-408-129, Thermo-Fisher Scientific). Paraformaldehyde (158127; Sigma-Aldrich). Phosphate buffered saline (PBS) (20-134; Apex Bioresearch Products). Sodium azide (NaN_3_) (S2002, Sigma-Aldrich). Tris Base (T8600, US Biological Sciences, Salem, MA). Tween-20 (P1379, Sigma-Aldrich).

### Rat Carotid Artery Balloon Injury

The rat carotid balloon injury model was first described by Clowes et al^4^. Since then it has been extensively described^31^. Shears et al. first described the modified, pressure-controlled segmental injury used in this study^32^. Adult male rats (n = 7 for LSFM, n = 10 for histology) aged 12-14 weeks were used for surgery. Rats were anesthetized with inhaled isoflurane (0.5–2%). Atropine (0.1 mg/kg) was administered subcutaneously (SC) to reduce airway secretions and Carprofen (5 mg/Kg) was administered SC for pain management at the time of surgery. After a sterile prep and midline neck incision, the left common, internal, and external carotid arteries were dissected and the internal and common carotid arteries were temporarily occluded using micro serrefine clamps (18055-05, Fine Science Tools, Foster City, CA). A No. 2 French Fogarty balloon catheter (Edwards Lifesciences, Irvine, CA) was inserted through an arteriotomy in the external carotid artery, and advanced into the common carotid artery. The balloon was inflated to 5 atmospheres of pressure for 5 min to create a uniform injury. After 5 minutes, the balloon was deflated and removed, the external carotid artery was ligated, and blood flow was restored. The incision site was sutured and rats were housed in a temporary cage under a heat lamp until consciousness was restored, then moved to their home cage. Rats were euthanized by anesthesia overdose followed by bilateral thoracotomy 2 weeks after injury.

### Rat Carotid Artery Harvest for LSFM

Both carotid arteries were harvested after in situ perfusion-fixation with 200 mL of 1X PBS followed by cold 4% paraformaldehyde. Arteries were harvested so that a portion of the bifurcation was included, as this helps with the sample orientation during imaging. Arteries were placed in 4% paraformaldehyde at 4°C overnight then moved to a 1.5 mL microcentrifuge tube with PBS + 0.02% NaN_3_ for 24 hours.

### Mouse Carotid Ligation

C57BL/6 mice were crossed with NOD mice (The Jackson Laboratory, Bar Harbor, ME) and used in this study. Mice (n = 3-6 for LSFM, n = 12 for histology) aged 12-14 weeks were anesthetized with inhaled isoflurane (1%-3%). Neck fur was removed with a depilatory agent, the field prepared with a 10% povidone-iodine solution, an incision was made slightly left of midline, and the left carotid artery just proximal to the bifurcation was exposed. Novafil™ polybutester (7-0) was used to ligate the carotid artery and close the incision. Mice were administered acetaminophen in drinking water (35mg/ml) for 4 days. Mice were euthanized 4 weeks after surgery under continuous isoflurane anesthesia (1.5%-3%) by cardiac puncture exsanguination with a 23g needle.

### Mouse Carotid Artery Harvest for LSFM

Four weeks after ligation, carotid arteries were harvested after in situ perfusion-fixation with 2-4% paraformaldehyde (PFA) in PBS. Arteries were fixed in 2-4% PFA at 4°C overnight and then moved to PBS + 0.02% NaN_3_ for 24 hours.

### Carotid Artery Staining and Processing for LSFM

Workflow is outlined in **Figure 1**. Arteries were blocked in IHC-Tek diluent overnight in a 37°C incubator. Rat arteries were probed with primary rabbit anti-CD31 (ab28364, Abcam) antibody diluted 1:500 in IHC-Tek diluent for 3 days at 37°C, washed four times over 24 hours in 1X PBS + 0.2% Tween-20 (PTwH wash buffer) at room temperature (RT), and probed with goat anti-rabbit Alexa Fluor 647 IgG secondary antibody (A-21245, Thermo-Fisher Scientific) diluted 1:500 in IHC-Tek diluent for 2 days at 37°C. Mouse arteries were stained following the same protocol with rabbit anti-CD31 (ab28364, Abcam) and probed with donkey anti-rabbit Alexa Fluor 568 IgG secondary antibody (A10042, Thermo-Fisher Scientific). For multiplex analysis mouse arteries were stained for CD31 in tandem with CD68 or VE-Cadherin. For the CD68 samples: 1:500 dilution of rabbit anti-CD31 and mouse anti-CD68 (MCA341GA, Bio-Rad, Hercules, CA) followed by a 1:500 dilution of Alexa Fluor 647 and goat anti-mouse Alexa Fluor 555 (A-21425, Thermo-Fisher Scientific) as described above. For the VE-Cadherin samples: 1:500 dilution of rabbit anti-CD31 and 1:300 dilution of goat anti-VE-Cadherin (AF-1002, R&D, Minneapolis, MN) followed by a 1:500 dilution of Alexa Fluor 568 and donkey anti-goat Alexa Fluor 633 (A-21082, Thermo-Fisher Scientific). Arteries were again washed in PTwH wash buffer over 24 hours at RT. After this wash, arteries were stored in PBS + 0.02% NaN_3_ in the dark at 4°C up to three weeks.

**Figure 1.**
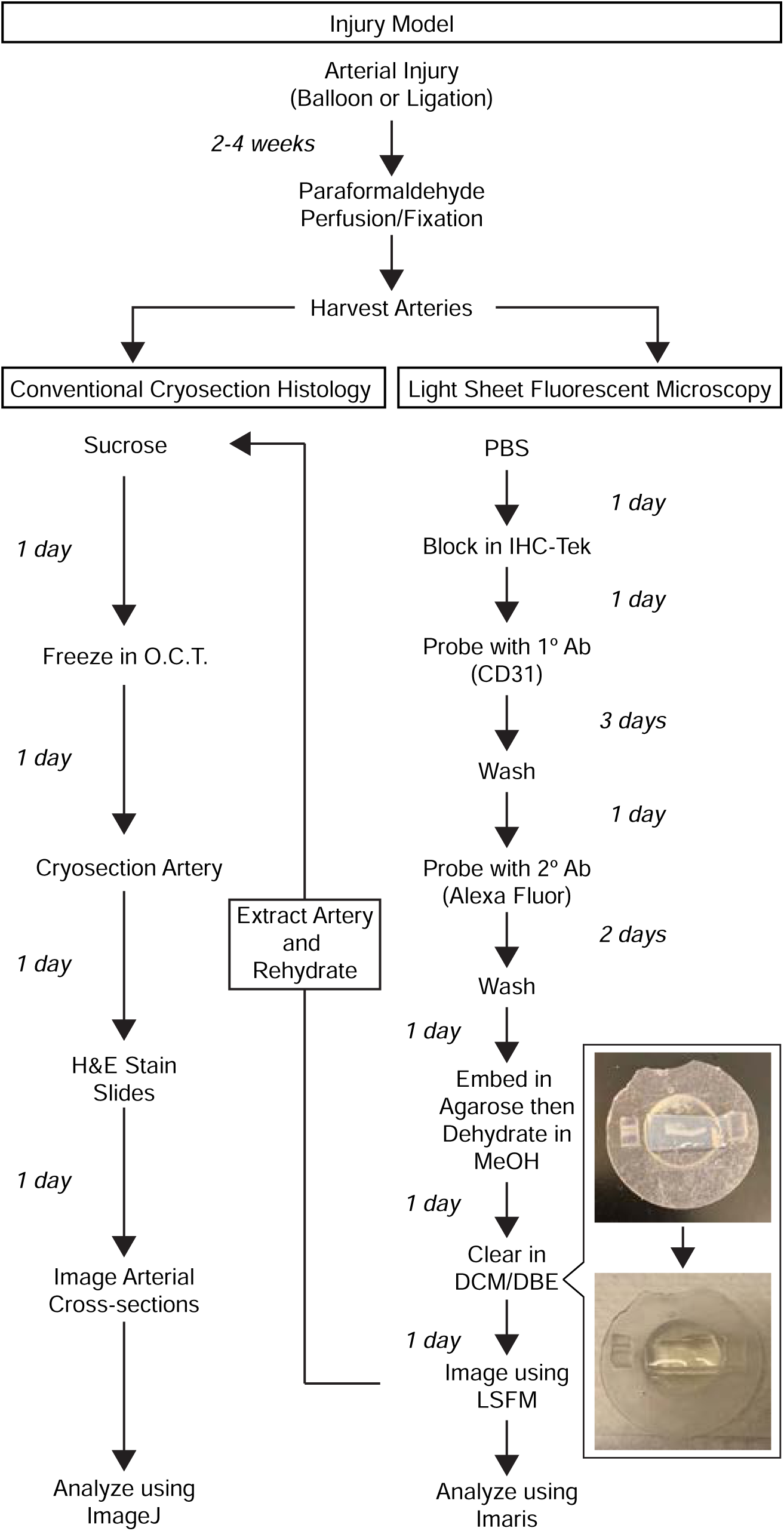
Sample processing workflow comparison for H&E or LSFM analysis. Arteries processed for LSFM imaging are able to be rehydrated and cryosectioned for H&E analysis.

Next arteries were embedded in 1% agarose (20-102GP, Apex Bioresearch Products, San Diego, CA) prepared in 1X TAE buffer. Agarose embedding was performed in a petri dish so that the final height of the agarose-artery block was 5 mm. To ensure success of the agarose embedding: 1) low-melt agarose should not be used as it is not compatible with iDISCO+; 2) prior to embedding, arteries should be warmed for over 1 hour in a 37°C incubator; 3) agarose temperature should be monitored using a thermometer and arteries should only be added once the agarose temperature is at 40°C to minimize heat damage. The final dimensions of the agarose-artery block that allow for proper sample mounting within the LSFM are 5 mm × 5 mm × 10-15 mm (W × H × L).

Arteries were then dehydrated in methanol (20% MeOH → 40% → 60% → 80% → 100% X2) for 1 hour at RT per dilution and then kept overnight at RT. Arteries were cleared the next day in glass scintillation vials (986492, Wheaton, Millville, NJ) on a plate shaker at RT. Arteries were exposed to a 33% methanol - 66% DCM solution for 3 hours, then 100% DCM for 15 minutes followed by a second round of 100% DCM for 15 minutes. Lastly, arteries were cleared in 100% DBE overnight. Arteries can be stored in the DBE indefinitely until imaging.

### Light Sheet Fluorescence Microscopy Imaging

Imaging was performed using a LaVision BioTec Ultramicroscope II^33^ equipped with zoom body optics, an Andor Neo sCMOS camera, an Olympus MVPLAPO 2X/0.5 objective and a 5.7 mm working distance corrected dipping cap (total magnifications ranging from 1.3X to 12.6X with variable zoom). Cleared agarose-artery blocks were mounted in a sample holder and submerged in a 100% DBE reservoir. Arteries were imaged at 1.6X mag (0.8X zoom), using the three light sheet configuration from a single side, with the horizontal focus centered in the middle of the field of view, an NA of 0.043 (beam waist at horizontal focus = 17 µm), and a light sheet width of 60-100% (adjusted depending on artery length to ensure even illumination in the Y axis). Pixel size was 3.9 µm and spacing of Z slices was 5 µm. Up to three channels were imaged per species: autofluorescence with 488 nm laser excitation and a Chroma ET525/50m emission filter was used for both species; mouse CD31-AF568 or CD68-AF555 stained channel imaged using 561 nm laser excitation and a Chroma ET600/50m emission filter; rat CD31-AF647 or mouse VE-cadherin-AF633 stained channel imaged using 647 nm laser excitation and a Chroma ET690/50m emission filter. The chromatic correction module on the instrument was used to ensure both channels were in focus. To reduce bleaching, while setting up the parameters for image acquisition the sample was viewed with low laser power and long exposures; this was inverted when acquiring data to maximize acquisition speed.

### Analysis Terminology

Internal elastic lamina (IEL) corresponds to the inner-most lamina, facing the lumen. External elastic lamina (EEL) corresponds to the outer-most lamina. As described below, an analysis tool is used to obtain the radius from the artery center to the edge of the lumen, touching the intima, or to either lamina. The change in radius (Δr) corresponds to the difference between the radius to the IEL surface edge and the radius to the lumen surface edge. This Δr is used to identify the region of hyperplasia though an automated, unbiased approach.

### Image Rendering and Analysis

Workflow is outlined in **Figure 2**. LSFM-saved TIFF files (Imspector Pro v5.1.350) were converted to an Imaris (Bitplane, Oxford Instruments) file using Imaris File Converter software (v9.3.1). Acquired LSFM images were first analyzed using the two-dimensional slice function and snapshots were obtained of cross-sections in the x-y, x-z, and y-z axis. The file was cropped to only include the artery to reduce file size and expedite all further steps. The Surfaces tool created three separate surfaces representing the artery lumen, IEL, and EEL. Each surface was created by semi-automatically tracing the appropriate landmark using the 647- or 561-channel for the CD31-stained intima, which would represent the lumen volume, and the 488-channel for both elastic laminas. Fluorescence captured in the 488-channel from the IEL was used as the spatial landmark for generating the IEL surface, while fluorescence from the EEL in the 488-channel was used for the EEL surface. For the rat model, contours were created using the isoline mode at 5% density along the artery every 15 μm in the x-z plane starting at the bifurcation until the end of the sample. For the mouse model, 15-20% density was used with traces every 10 μm. For some optical slices the semi-automatic tracing failed to identify the landmark. In these cases, a manual trace was performed. The starting slice position at the bifurcation and final slice position were noted for the first rendered surface, to ensure that all three rendered surfaces visualized the same arterial section. This was first performed on the healthy artery for each animal, and then on the injured artery.

**Figure 2.**
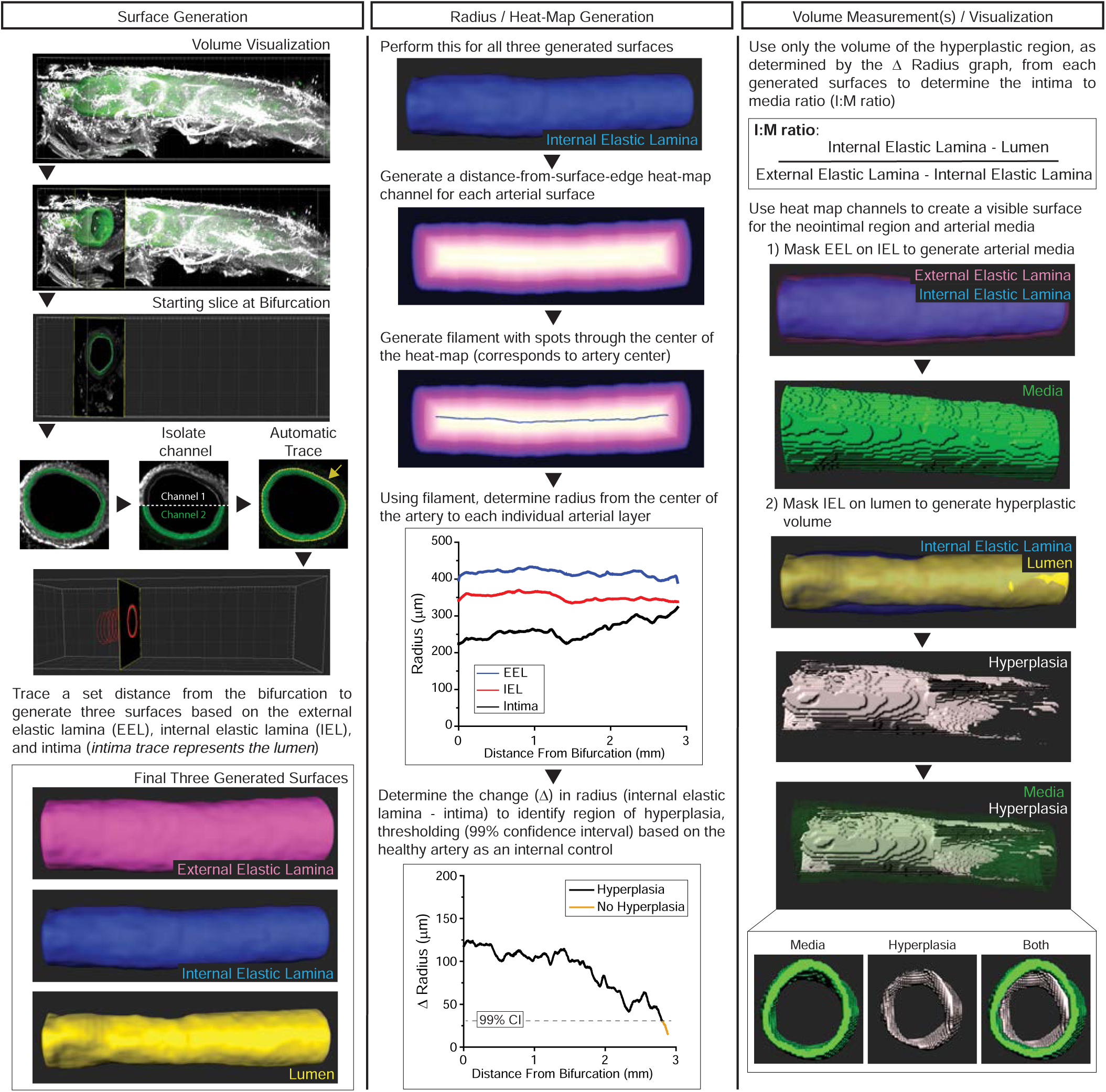
Analysis workflow in Bitplane Imaris. Rat carotid artery used as the representative sample. Video 2 overlays the three surfaces with the fluorescent volumetric rendering (https://youtu.be/LoAIApmV22o).

Using the IEL surface, a filament was generated through the center of the artery. A Distance Transformation was performed inside the IEL surface to generate a heat-map along the artery from the artery center to the surface edge. This heat-map is presented as a new channel whose intensity corresponds to the distance in µm, from the surface edge. This was replicated for the other surfaces, and channels were named “Distance to ‘respective surface’”. Next, the Filament tool, using the Line style with auto-center and auto-depth activated, was used on the IEL heat-map channel to generate a filament through the center of the heat-map, corresponding to the artery center. Then the generated filament was converted using a Filament to Spots XTension (provided by Matthew Gastinger, Bitplane), which automatically placed an individual spot roughly every 4 μm along the artery. Using these Spots, the radius along the artery from the center of the artery to the surface edge of the lumen, IEL, and EEL was obtained. The average intensity at each spot (which corresponds to the radius, in µm) and the spot position were exported to Microsoft Excel. Cumulative distance along the filament (i.e. the center of the artery) was calculated using a Pythagorean formula for Euclidean distance between adjacent spots. Up to 50 spots from the lateral edges of the filament were removed because they measured the distance to the lateral region of each surface, which is inconsistent with the rest of the spot measurements along the filament. A plot was generated of the radius to each surface along the artery. The change in radius (Δr = radius to IEL surface edge – radius to lumen surface edge) was obtained to identify regions of hyperplasia. To control for potential sample processing artifacts and to provide baseline parameters, we used the contralateral, uninjured carotid artery as a reference per animal. In the healthy artery the intima is a single monolayer of endothelial cells, thus the Δr centers close to zero. Using the average Δr for the uninjured artery, we estimate the 99% confidence interval (CI) and define any Δr above this threshold as indicative of neointimal hyperplasia in the animal’s contralateral injured artery. In the injured artery, the spot positions above the 99% CI were noted. Using the identified starting and ending spot positions, all three surfaces were cropped in Imaris, which generated three new surfaces exclusively in the region with neointimal hyperplasia. Using the newly cropped surfaces, the volume of each respective surface was exported to Excel and the intima-to-media (I:M) ratio ((IEL volume - lumen volume) / (EEL volume – IEL volume)) was obtained.

To visualize the hyperplasia and media (as shown in the right panel of **Figure 2** or in **Figure 3F**) we generated new surfaces using a combination of the Imaris Surface tool and masking of the distance transformation channels. We defined the media volume as the space between the IEL and EEL surfaces, and the hyperplasia as the space between the lumen and the IEL surface.

**Figure 3.**
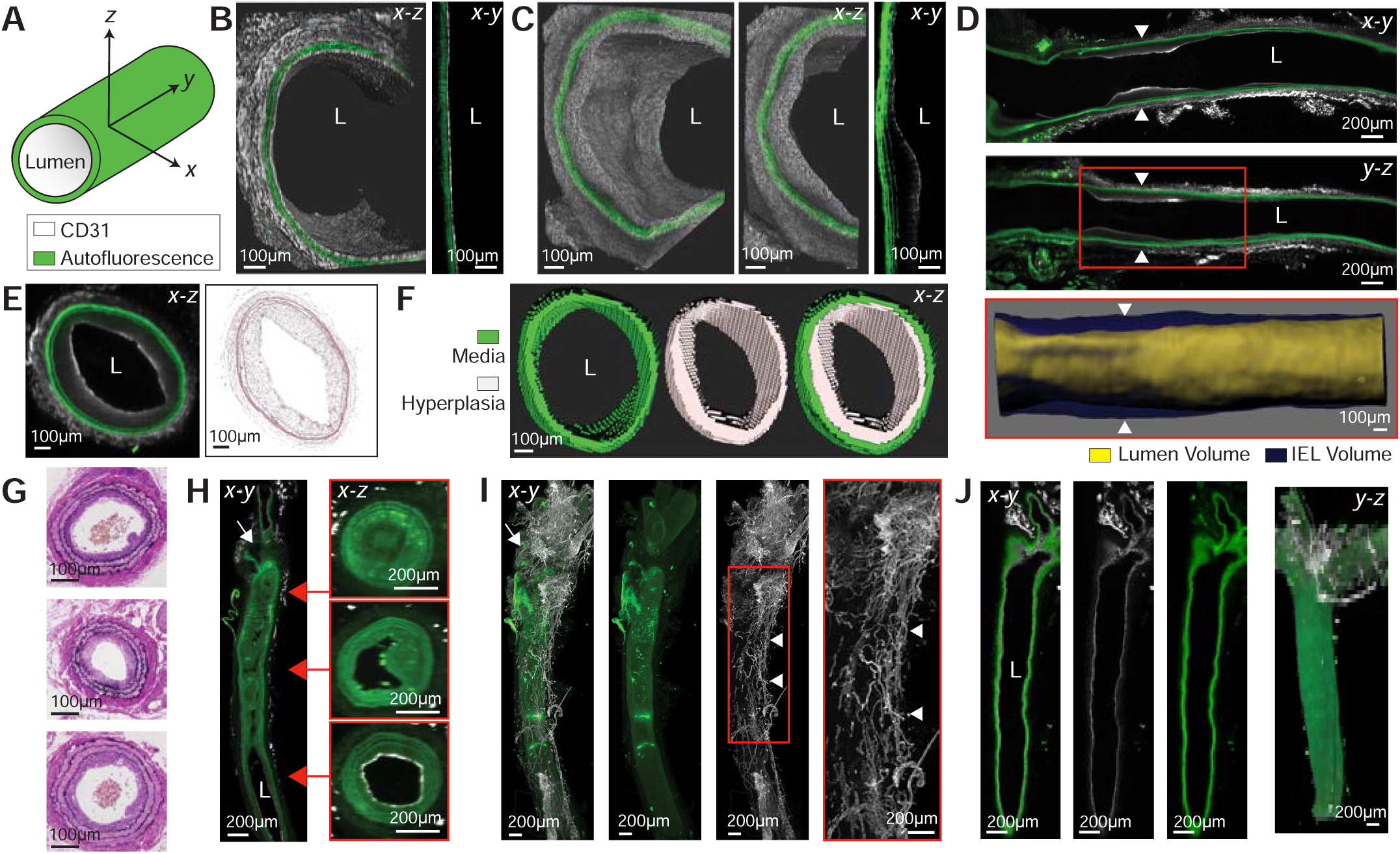
Demonstration of LSFM imaging for rat and mouse injury models. **(A)** Vessel orientation and figure image colors, unless otherwise stated. **(B)** Volumetric rendering and slice view of healthy rat artery (L = lumen). **(C)** Representative balloon-injured rat artery. **(D)** Representative slice views of an injured rat artery (arrowheads = hyperplasic region). White box indicates region of the surface rendering of the internal elastic lamina (IEL) and lumen volumes. **(E)** Representative cross-section in x-z axis from arrowheads in (D) with H&E-stained image of the same artery after rehydration and cryosectioning. **(F)** Cross-section of surface rendering of the representative rat (from (D) and (E)) arterial media, hyperplasia, and both surfaces overlaid. **(G)** Representative H&E-stained cross-sections of ligated mouse artery processed exclusively for histology. **(H)** Longitudinal and cross-sectional slice rendering of representative ligated mouse carotid artery (white arrow = ligation site at bifurcation) **(I)** Representative volume rendering of ligated mouse carotid artery depicting the periadventitial neovascular plexus (white arrow = ligation site at bifurcation, arrowheads = neo-vascular plexus). **(J)** Slice and volumetric rendering of representative healthy mouse artery.

### Carotid Artery Histological Processing and Histology

Rat or mouse carotid arteries were harvested after in situ perfusion-fixation with PBS and cold 2% paraformaldehyde. Arteries were placed in 2% paraformaldehyde for 1h at 4°C followed by 30% sucrose overnight at 4°C. Arteries were snap-frozen in O.C.T. (4583; Tissue-Tek, Torrance, CA) and stored at −80°C. 5 μm sections were cut throughout the entire common carotid artery for staining.

After imaging for LSFM, the agarose-embedded artery was rehydrated in PBS + 0.02% NaN_3_ for 3 days in a 40°C water bath. Artery was manually extracted and soaked overnight in 30% sucrose followed by cryosectioning and staining.

Arteries were hematoxylin & eosin (H&E) stained for morphological analysis and the LSFM-processed, rehydrated artery cross-sections were also stained for immunofluorescence. Arterial cross-sections on room temperature slides were fixed with ice-cold acetone for 5 min then permeabilized with 0.3% Triton X-100 (X100; Sigma-Aldrich) for probing with 1:100 primary anti-alpha-smooth muscle actin (α-SMA) antibody (Rabbit, ab5694, Abcam, Cambridge, UK) diluted in IHC-Tek diluent (1W-1000; IHC World, Woodstock, MD) followed by 1:500 dilution of goat anti-rabbit Alexa Fluor 555 IgG secondary antibody (A-21429, Thermo-Fisher Scientific) and 1:500 DAPI diluted in PBS. Slides were rinsed in distilled water and coverslips were attached using ProLong Gold antifade reagent (P36930, Invitrogen). Slides were imaged on the day after staining using a Zeiss Axio Imager.A2 microscope (Oberkochen, Germany). Samples were illuminated by the X-Cite 120 LED Boost (Excelitas Technologies, Waltham, MA) laser with DAPI (385 nm), GFP (470 nm) or Cy3 (560 nm) excitation filters. Images were taken on a 5X objective (Zeiss Plan Apochromat 5x.0,16 440620 (2)) for quantification of rat arteries or a 10X objective (Zeiss Plan Neofluar 10x/0.30 440330) for mouse arteries. 20X objective (Zeiss Plan Apochromat 20x/0.8 ∞/0.17) images of the rat arteries were included for supplemental figures. Images were captured using an AxioCam HRm (Carl Zeiss MicroImaging GmbH, 426511-9901-000 10-33 VDC 5W) camera.

### Histological and Optical Slice Analysis

All images were quantified using ImageJ software (https://imagej.nih.gov/ij/). Intima-to-media (I:M) ratio per animal was determined by dividing the area of neointimal hyperplasia (IEL-lumen area) by the medial area (EEL-IEL area) per optical or histological slice per animal. 6-10 discontinuous arterial cross-sections were analyzed for the injured rat carotid arteries comparison to LSFM volume and optical slice I:M ratio. 3-4 discontinuous arterial cross-sections were analyzed for the healthy rat or mouse carotid artery arterial shrinkage analysis.

Virtual snapshots for the mouse or rat were taken at a constant zoom for each animal per species. Up to 10 virtual snapshots of the injured rat artery in the x-z axis were taken every 300 µm for analysis around the peak slice of injury. Area and perimeter values were obtained of the external elastic lamina, internal elastic lamina (IEL), and lumen. Up to 10 virtual snapshots of mouse arteries were analyzed every 100-200 µm along the carotid artery. Then 3-10 slices, as defined in the figures, were averaged per animal.

### Statistics

For all multiple group analyses, a one-way ANOVA with Tukey’s correction was performed in Origin software. Two group analysis was performed using a student’s t-test, and paired-group t-test when appropriate, in Origin software. One group comparison to a reference value was analyzed with a one sample t-test. Precision was assessed using the coefficient of variation (CV%) for the 3-D volume to 2-D areal analysis. Rat arterial remodeling was analyzed with SAS statistical software (SAS Institute). The partial correlation coefficient (ρ) between neointimal thickness and vessel radius was estimated using a mixed model^34^, controlling for subject variation and in repeated measures along the length of the artery. The confidence interval and p-value for the mixed model correlation estimator was determined with a normal approximation using the delta method^35^. SAS Code provided in the online supplement.

## III. Results

The aim of this manuscript is to validate LSFM for analysis of preclinical restenosis models. The combination of iDISCO+ with LSFM is ideally suited to image injured arteries, generating datasets to measure multiple aspects of the arterial injury response efficiently while minimizing bias. This technique generates optical sections of the transparent sample that can then be processed for analysis (**Videos 1-3** –see legend for hyperlink to videos–). A comparison of the workflow for histological and LSFM processing is provided in **Figure 1**, while the analysis workflow utilizing Bitplane Imaris software is depicted in **Figure 2**.

### Imaging of injured murine arteries

First, we set out to image and analyze the rat carotid balloon injury model. LSFM imaging of injured and contralateral uninjured arteries clearly allows for identification of the three layers of the arterial anatomy: intima, media (including the individual lamellae) and adventitia (**Fig 3 B-E)**. A close-up volume rendering and x-y plane optical slice of the healthy (**Fig. 3 B**) and injured (**Fig. 3 C**) rat arteries shows a clear CD31^+^ endothelial layer and clear morphological differences on the luminal side between the uninjured and injured arteries. To identify the area of neointimal hyperplasia we present two longitudinal views of the rat injured artery from the bifurcation through 5 mm along the common carotid (**Fig. 3 D**). A surface rendering of the hyperplastic region is presented, delineated by the red box (**Fig. 3 D**). This rendering is performed for the hyperplastic region of all arteries to measure vessel stenosis, and an overlay of this rendering onto the whole artery is shown in **Video 2** (https://youtu.be/LoAIApmV22o). To compare the 2D images obtained through LSFM with those obtained through H&E staining and histological imaging, **Figure 3 E** presents a transverse (x-z) cross-section of the artery in **Figure 3 D**, side-by-side with an H&E-stained cross-section of the same sample after it was rehydrated, cryo-sectioned, and stained. **Figure 3 F** depicts a surface rendering of the hyperplastic lesion and the media of the injured rat carotid artery, the volumes of these regions are used to analyze stenosis through volumetric I:M ratio. To show the intra- and inter-subject variability of the model, representative slices in the longitudinal x-y and y-z planes and determination of the region of hyperplasia of each rat artery used for analysis are shown in **Sup. Figure 1**.

Even though the rat carotid balloon injury is a commonly used model, the availability of many genetically modified mouse strains makes mouse models of arterial injury very important tools in vascular biology. Hence, it was of interest to determine if our methodological approach is suitable for arteries that are ten times smaller than those of the rat. Therefore, we assessed the ability of our imaging and analysis methodology in the mouse carotid ligation model (**Fig. 3 G-J)**. We compare the results obtained through LSFM with those obtained by histological imaging of H&E-stained sections. We show representative H&E cross-sections from three ligated mouse arteries processed exclusively for histology (**Fig. 3 G**) next to longitudinal (x-y) and transverse (x-z) slice views of a representative ligated mouse artery imaged through LSFM (**Fig. 3 H**). Notably, the transverse cross-sections corresponding to the different regions of the artery highlight the variability in neointimal hyperplasia development within a single artery. The intra- and inter-subject variability can be observed in **Sup. Fig. 2**, which shows representative slices of each mouse artery used for analysis. Specific to the mouse ligation model, we observed a CD31^+^ periadventitial neovascular plexus around the ligation and along the injured artery (**Fig. 3 I**). This CD31^+^ plexus was not present in the contralateral uninjured artery (**Fig. 3 J**). This neovascularization was consistent for all ligated arteries (**Sup. Fig. 2**), including arteries with low (<0.1) I:M ratios, and presents a phenomenon that has not been described in rodents by classic histological approaches.

Inflammation is a critical characteristic of the arterial injury response, and macrophages are known to be present at the site of injury. Moreover, inflammatory macrophages can drive angiogenesis^36^. Therefore, it was of interest to assess the presence of macrophages in injured arteries while assessing the multiplexing capabilities of our methodology. To do this, some arteries were stained for CD68, a commonly used macrophage marker, together with CD31, prior to LSFM imaging. **Figure 4** shows presence of the neovascular plexus and macrophages in the adventitia of the injured carotids evidenced by the CD31^+^ staining and CD68^+^ puncta. To further assess the multiplexing capabilities of the technique, some uninjured arteries were stained for VE-cadherin, an endothelial cell adhesion marker, and CD31. **Sup. Figure 3** shows optical slices and optical volumes of uninjured mouse carotids showing the traditional VE-cadherin^+^ cobblestone structure of intact endothelium and the CD31^+^ cells. These experiments show that multiplexing in mouse arteries is possible using this methodology, further expanding the capabilities of LSFM in preclinical arterial injury models.

**Figure 4.**
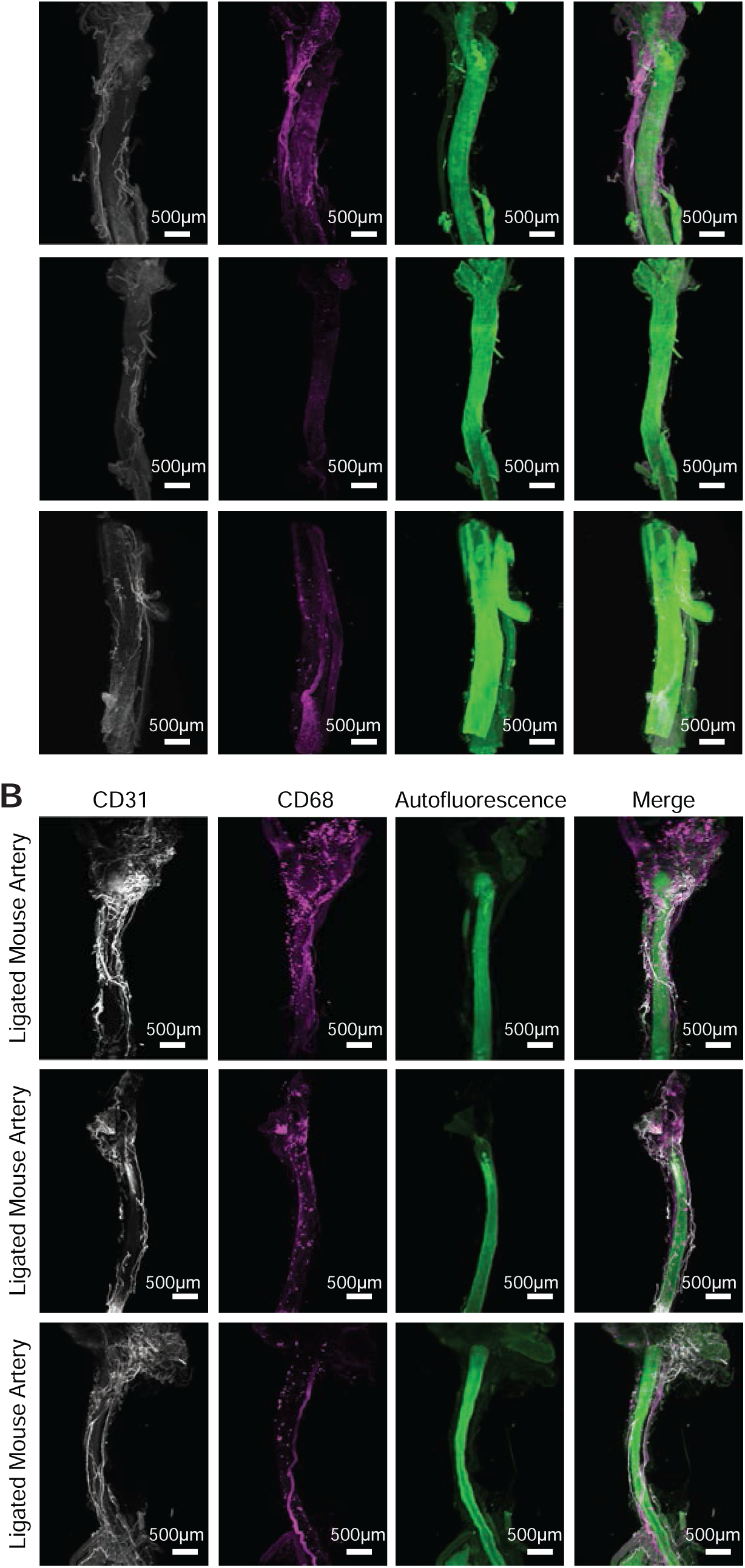
Macrophage presence in the adventitia of injured mouse carotids. Uninjured and ligated mouse carotid arteries stained for CD31 and CD68 show presence of CD68^+^ puncta only in the ligated arteries around the ligation site and periadventitially next to the CD31^+^ neovascular plexus (n = 3, white = CD31, magenta = CD68, green = autofluorescence). **(A)** 3D rendering healthy mouse carotid arteries. **(B)** 3D rendering healthy mouse carotid arteries.

### Quantification of arterial stenosis

Next, we compared the quantitative capabilities of our methodology with those of traditional histology. We used the intima-to-media (I:M) ratio for both injury models to benchmark the LSFM volumetric analysis approach to classic histology. To do this, we compared the volumetric I:M ratio to the H&E I:M ratio of a separate cohort of age-matched animals. The results (**Fig. 5 A-B**) showed comparable average I:M ratios with no significant difference for both rat (LSFM = 0.77, H&E = 0.62) and mouse (LSFM = 0.79, H&E = 0.63), thus validating our methodology. However, the coefficient of variation (CV%) of the estimates was lower for the LSFM volumetric approach compared to traditional H&E histology (Rat = 28% vs 41%, Mouse = 79% vs 127%; **Sup. Table 1**) suggesting increased precision. Additionally, LSFM imaging allows for optical slicing of the artery as thin as 3.2 µm along the x-z axis. Thus, per animal model, we took equidistant optical slices for each injured artery and analyzed the slices using the classic 2-D area trace. We evaluated the utility of analyzing 3, 5, and 10 optical slices per animal, as these are within the commonly reported number of cross-sections analyzed in the literature (**Table 1**). We then compared these values to the volumetric approach and observed no significant difference between the volumetric and classical analyses (**Sup. Fig. 4**). However, for the rat, the more optical slices used, the more precise the I:M estimate, as illustrated by the CV% (**Sup. Table 1**). For the mouse ligation model, any slice number used for analysis had a CV% similar to the volume approach, and almost half of that for the H&E-analyzed samples (optical slice = 65-69% vs. H&E = 127%, **Sup. Table 1**). The lower precision of the estimates in the mouse carotid ligation model can be explained by the higher intra- and inter-subject variability of the model (**Sup. Fig. 1 and 2**) Our results suggest that analysis using volumetric renderings produces similar estimates to classical histological approaches with greater precision, while allowing for optical sectioning.

**Table 1.**
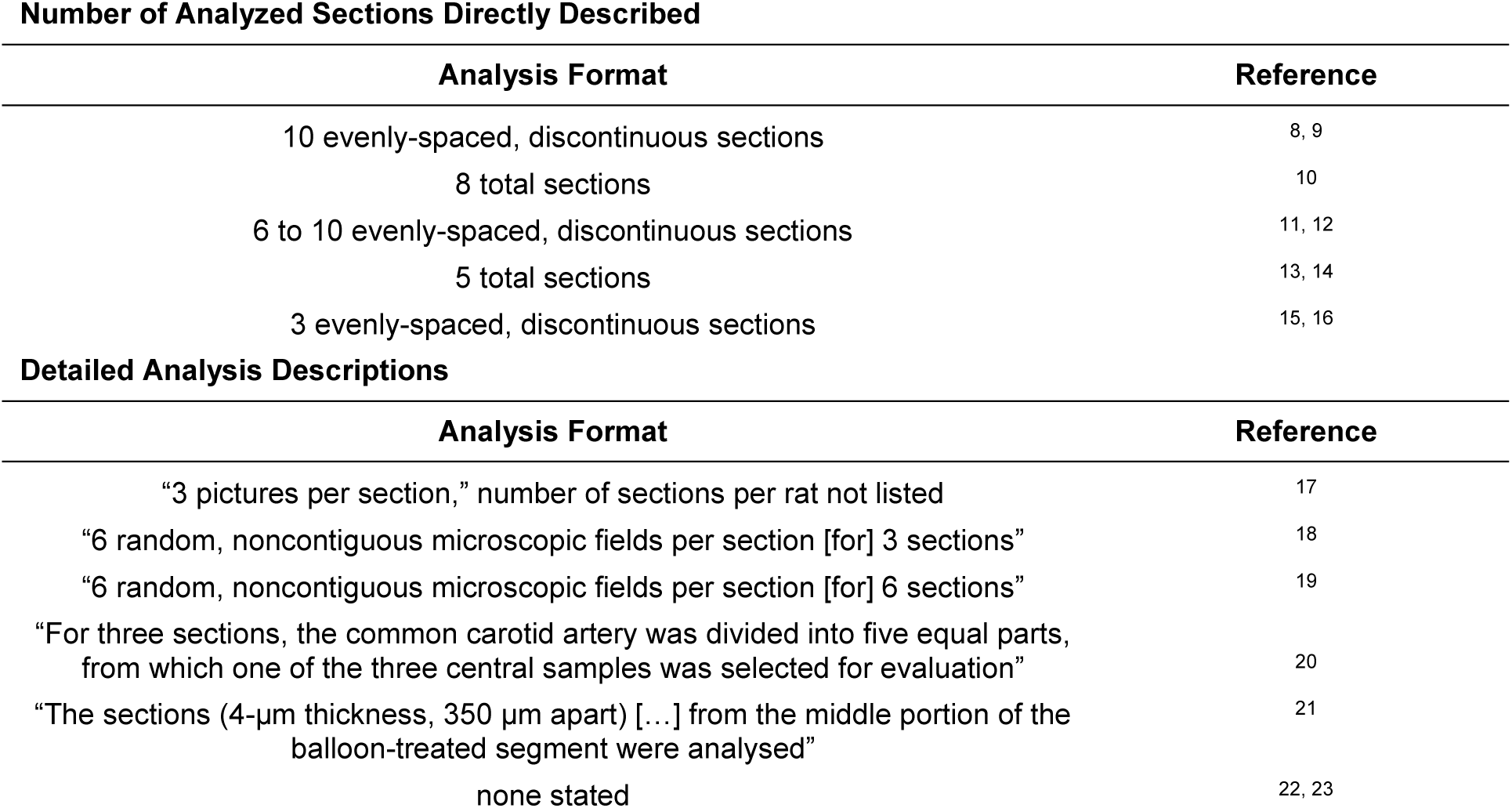
Neointimal hyperplasia analysis in rat balloon injury models according to published method sections.

**Figure 5.**
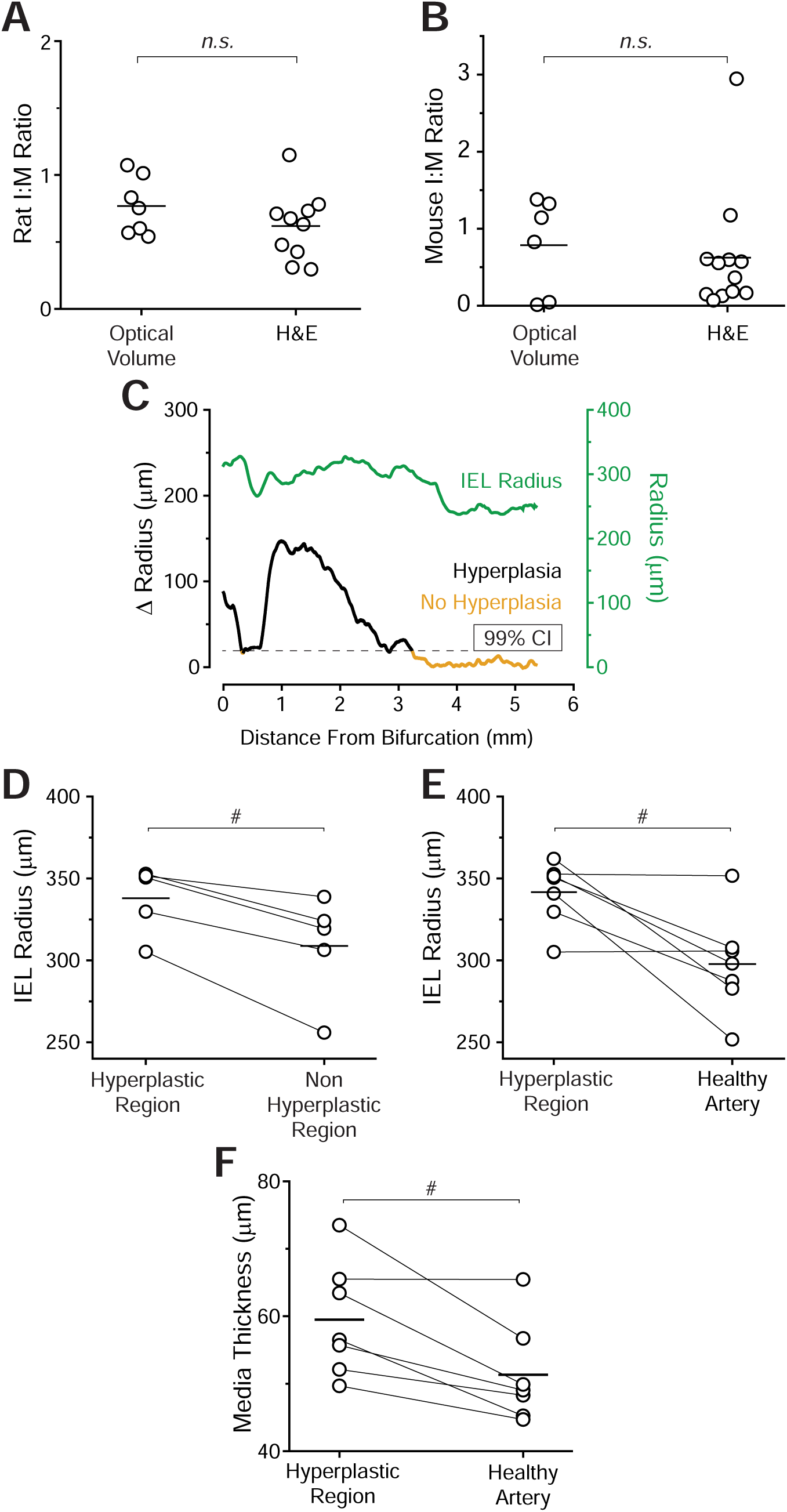
Demonstration of LSFM quantification. **(A)** Rat intima-to-media (I:M) ratio using optical 3-D volume compared to a separate cohort of injured rat arteries harvested directly for histological processing (LSFM n = 7, H&E n = 10, n.s. = no significance, student’s t-test). **(B)** Mouse I:M ratio using optical volume compared to a separate cohort of ligated mouse arteries harvested directly for histological processing (LSFM n = 6, H&E n = 12). **(C)** Plot of injured rat artery IEL radius and change (Δ) in radius between the lumen and the IEL along the artery center (dataset shown in Fig. 3 D). 99% confidence interval (CI) defines the threshold at which the Δ Radius is indicative of neointimal hyperplasia formation. 99% CI determined using the average Δ Radius along the contralateral healthy rat artery (Sup. Fig. 1). **(D-E)** Average IEL radii per rat along their hyperplasic region compared to the non-hyperplastic region of the injured artery or to their contralateral healthy artery (^#^p<0.05, paired t-test). **(F)** Plot of medial thickness comparing the hyperplastic region of each injured artery to the respective contralateral healthy control artery.

Importantly, the use of the Filament analysis tool allowed for unbiased identification of the hyperplastic region (**Fig. 2**). **Figure 5 C** shows a representative plot of the IEL radius overlaid with the change in radius (Δr = IEL radius – lumen radius). In the healthy artery the lumen and IEL radii are practically identical, thus the Δr centers close to zero. Meanwhile, in the injured artery there is a dramatic increase in Δr.

### Quantification of remodeling

The ability to look at the whole artery led us to assess the quantification of vessel remodeling after arterial injury. Notably, the arterial filament allowed us to measure positive vessel remodeling within the hyperplastic region in the rat model. We compared the individual spots corresponding to the hyperplastic region along the filament to the non-hyperplastic region of the injured artery and to each rat’s respective contralateral healthy artery. Our results (**Fig. 5 D-E**) show a significant increase in IEL radius within the hyperplastic region (p=0.008 to non-hyperplastic regions within the same vessel, p=0.01 to healthy). Furthermore, intimal thickness (T) correlated positively with vessel radius (R) (ρ_TR_=0.58; 95% CI 0.38-0.80; p<0.0001) (**Sup. Table 2**) indicating compensatory remodeling in the identified region of neointimal hyperplasia. Additionally, constrictive remodeling was evidenced by an increase in medial thickness in injured vessels. The medial thickness of the hyperplastic segment was on average 8 µm thicker per rat compared to their contralateral healthy artery (p=0.01) (**Fig. 5 F**), but was not significantly different from the thickness of nearby uninjured arterial segments (p=0.512) (data not shown).

### Validation of morphological parameters after LSFM processing

Even though the iDISCO+ clearing method we used has been shown to preserve morphology and size of cleared samples^30^, it was important to verify that there was no arterial shrinkage or expansion in our experimental approach. Therefore, we compared the luminal surface area and perimeter, or EEL perimeter, between LSFM-processed and H&E-processed age-matched healthy arteries for both the rat (**Fig. 6 A-C**) and mouse models (**Fig. 6 D-F**). As noted in **Figure 1**, LSFM processing is reversible by removing the artery from the agarose block and rehydrating it. **Figure 7** shows representative α-smooth muscle actin immunofluorescent staining and H&E staining of a rehydrated artery. This verifies that arterial samples processed for LSFM imaging can be used to analyze other biological parameters *a posteriori*. To assess if the rehydration process affects arterial size, we compared the I:M ratio and percent occlusion obtained by the optical volume to the optical slice or rehydrated H&E slice analysis for a single rat artery. Equally spaced optical sections along the region of hyperplasia of one rat were analyzed for the I:M ratio, percent occlusion, and perimeter. After rehydration, cryosectioning, and H&E staining, equally spaced sections were quantified and compared to the results from the optical slices. No significant differences were found for any of the parameters assessed (**Sup. Fig. 5**). However, the precision of the optical slice measurement was still higher as evidenced by lower CV% (38% for the percent occlusion estimate using optical slices vs 52% using rehydrated and H&E-stained slides).

**Figure 6.**
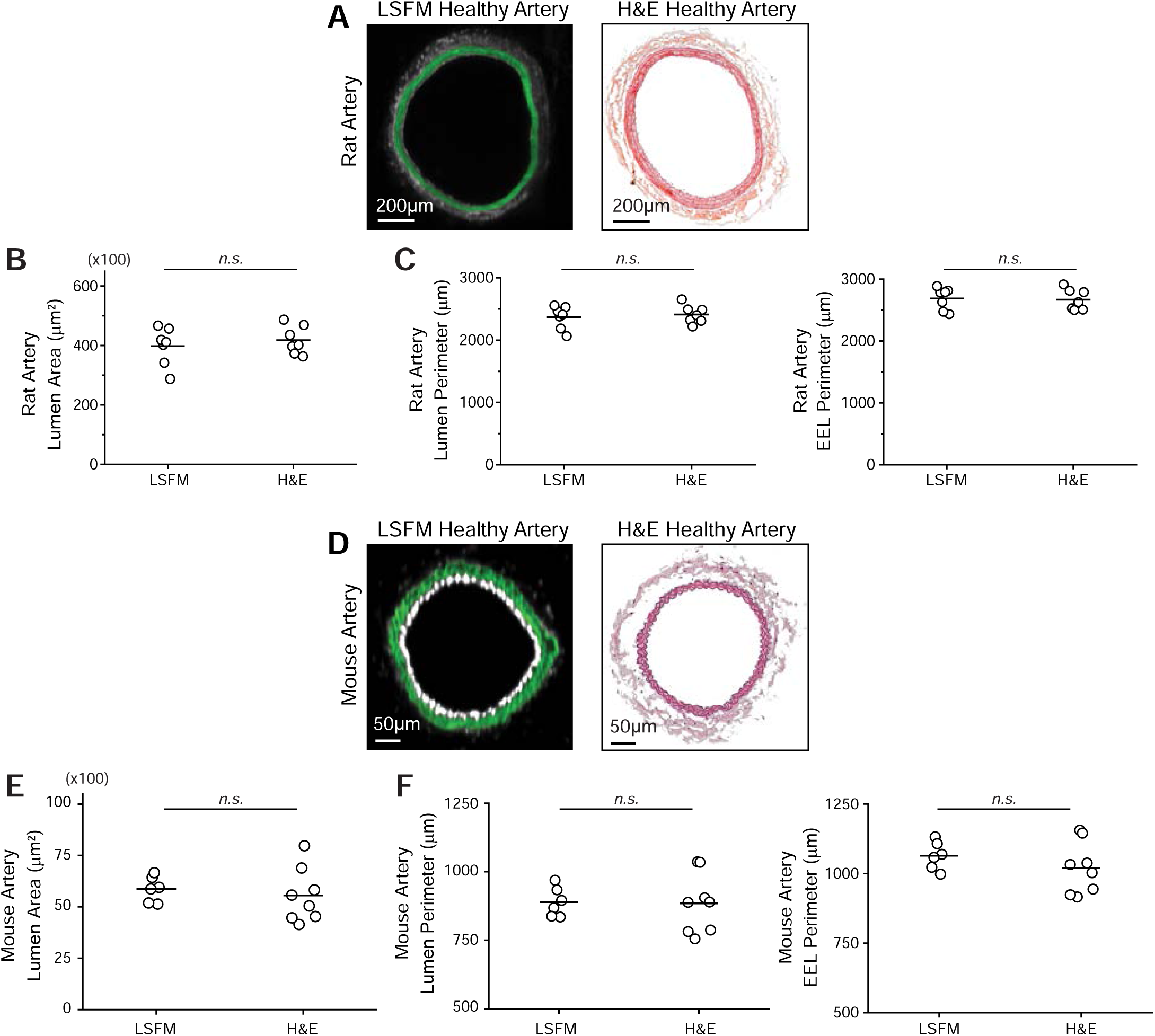
Artery processing for LSFM imaging does not cause arterial shrinkage. Comparison between healthy, uninjured arteries processed only for LSFM or only for hematoxylin & eosin (H&E) analysis. **(A)** Representative cross-section of healthy rat arteries. **(B)** Average luminal surface area per healthy rat artery (n = 7, n.s. = no significance, student’s t-test). **(C)** Average luminal and external elastic lamina (EEL) perimeter per healthy rat artery. **(D)** Representative cross-section of healthy mouse arteries. **(E)** Average luminal surface area per healthy mouse artery (LSFM n = 6, H&E n = 8). **(F)** Average luminal and EEL perimeter per healthy mouse artery.

**Figure 7.**
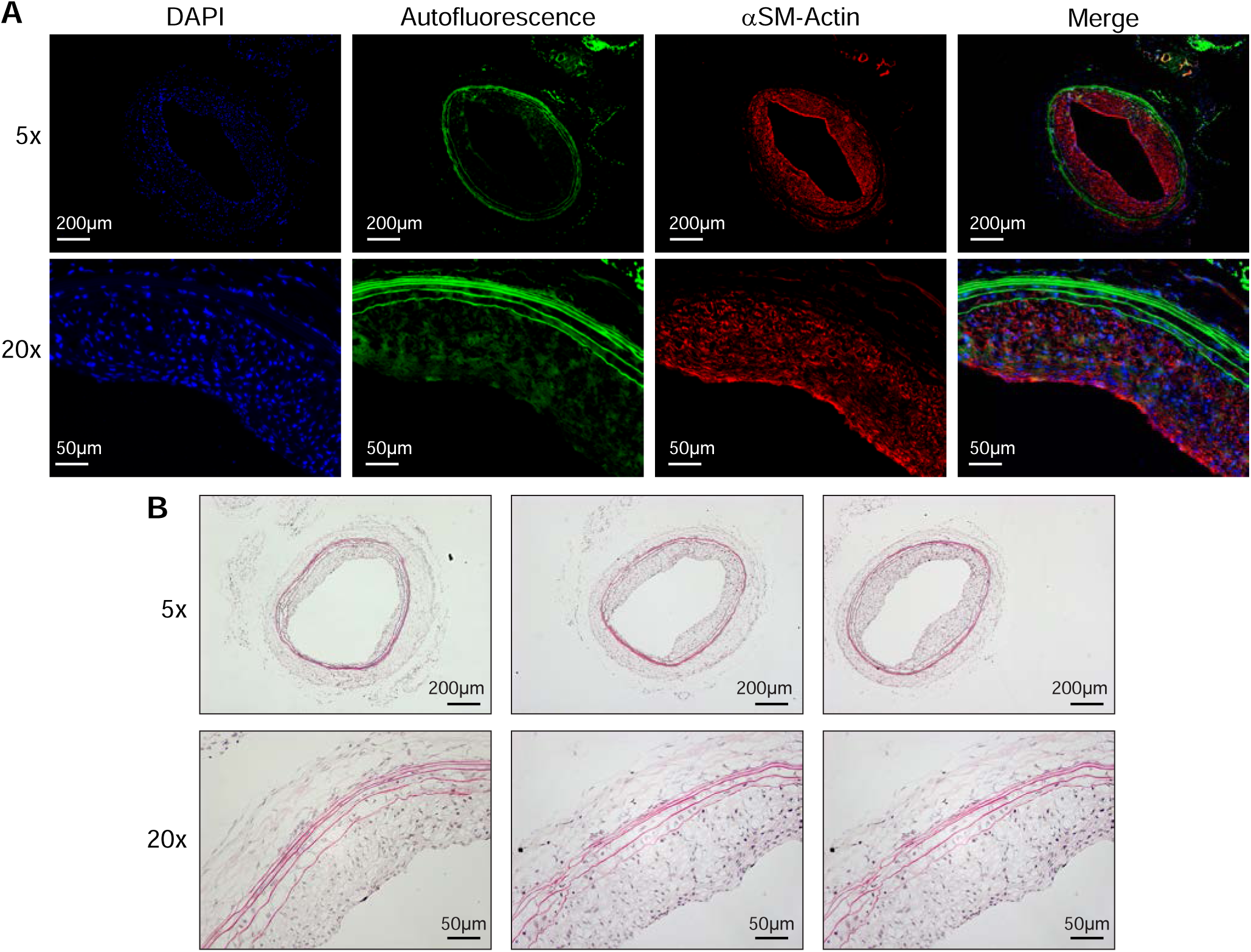
Histological staining of rehydrated, LSFM-processed rat carotid artery. **(A)** Immunofluorescent staining for alpha-smooth muscle (αSM) actin. **(B)** H&E staining of three representative arterial cross-sections.

## IV. Discussion

We describe a platform for analyzing vascular injury models in both mouse and rat arteries in 3-D. While the maximum resolution of the LSFM used herein is lower than that of alternative imaging techniques, it can precisely measure key parameters of arterial injury in both rat and mouse injury models without the need for physical sectioning. Further, the reconstructed LSFM images in 3-D allow for several approaches to injury quantification of the same sample. As such, this study shows that LSFM can be used to analyze neointimal hyperplasia formation and arterial remodeling after injury and outperforms traditional histology methods in quantitative capabilities and precision.

The advantages of utilizing 3D datasets and volumetric analysis compared to classical 2D analysis have been studied by Gorelashvili *et al.* in the bone marrow^27^. They showed that LSFM followed by 3D volumetric analysis of cells in the bone marrow reduces analysis bias and provides additional information to classically used techniques. Here we draw analogous conclusions regarding the power of this methodology when used to analyze vascular injury and subsequent stenosis. In the rat model we were able to obtain more precise estimates of stenosis and more detailed quantitative information about remodeling compared to histology. We compared vessel diameter and medial thickness in areas of hyperplasia to the contralateral artery and regions without hyperplasia within the same artery. Additionally, we were able to use a repeated measures design to establish a correlation between intimal thickness and vessel diameter. In the mouse model, we obtained more precise estimates of stenosis compared to histology, and we found periadventitial neovascularization of injured arteries.

Classic histological methods for analyzing neointimal hyperplasia development provide an incomplete picture as a select few cross-sections are used to inform the full extent of pathological changes that occur in the vessel. The classic approach introduces user bias in slide selection for analysis and is not standardized throughout the scientific literature (**Table 1**). Furthermore, an extensive literature review of preclinical cardiovascular studies revealed that methodological sources of bias compound the preexisting limitations of interpreting animal data in modelling human disease^37^. Thus, poor translation of preclinical models is in part due to the lack of rigor and reproducibility in preclinical study design and analysis. LSFM markedly improves the rigor of vascular disease assessment and is likely to improve reproducibility of preclinical findings. This is primarily because LSFM reduces user bias by imaging the entire intact artery, which can then be visualized in 3-D using Bitplane Imaris. Importantly, we show that tissue processing for LSFM imaging and analysis, and subsequent rehydration and cryosectioning, does not induce arterial shrinkage or expansion. Thus, the necessary parameters for quantifying vessel stenosis remain unaltered by processing. The obtained I:M ratio is comparable to that obtained by classic histology, but is more precise as evidence by the lower CV%. Furthermore, the Filament tool in Bitplane Imaris allows for unbiased identification of hyperplastic regions, and detection of compensatory remodeling and medial thickening throughout the artery. It is important to note that even though sample processing time is longer than traditional processing for histology (**Fig. 1**), most of it is not active researcher time. Intravenous injection labeling for CD31 is a possible alternative strategy to expedite the staining protocol. This approach has been shown to effectively label endothelial cells in the heart^38^ as well as in the kidney^39^. Additionally, it is also worth noting that there is a learning curve to become proficient with Bitplane Imaris analysis software. However, once enough proficiency is achieved, analysis time is not significantly longer than traditional histology. Importantly, unlike lengthy cryosectioning, analysis time allows for direct interaction with the data in a more intellectually engaging manner.

To our knowledge, the observation of the neovascular plexus around the adventitia of ligated mouse artery has not been previously reported. Kwon et al. showed that the adventitial vasa vasorum of pig coronaries increased after balloon injury. The neovascularization 28 days after injury was proportional to the degree of stenosis^40^. No such finding had been reported for murine models. The periadventitial CD68^+^ regions in injured arteries close to the neovascular plexus suggest that inflammation might contribute to the angiogenic process. However, the process leading to the development of the periadventitial neovascular plexus in the injured arteries grants further research that is beyond the scope of this work. Nonetheless, our observation exemplifies the potential LSFM has in driving biological discoveries in murine models of vascular injury.

Compared to confocal microscopy, LSFM has significantly reduced photobleaching. That said, photobleaching can still be a problem if, during imaging, a single optical section is exposed to laser light for a very long time. In that case, the resulting photobleaching generates a darker x-y plane in the sample, which shows up as a “break”; this makes semi-automatic tracing on vessels (**Fig. 2**) more difficult. This problem can be circumvented by minimizing exposure during sample setup and, post hoc, by manually tracing contours in the vessels if semi-automatic tracing does not work. Biological studies often involve the need to study different molecular markers at once. Hence, multiplexing capabilities are an important feature of powerful techniques. Multiplexing using LSFM has been successfully used to image kidney organoids^41^, the mouse brain^42^, and human neural aggregates^43^. Here we show the labeling of two different molecular markers at once to exemplify the multiplexing capabilities of the technique in studying vascular injury. We show concurrent staining for CD68 and CD31, and for VE-cadherin and CD31. The resolution of this technique is enough to show the characteristic morphology of the endothelium as evidenced by the cobblestone pattern of VE-cadherin staining. However, when comparing the resolution of this form of LSFM to confocal microscopy, LSFM does not allow visualization of subcellular details; being able to rehydrate the tissue and do high-resolution confocal microscopy on sections allows multiple levels of detail to be probed, from the tissue-wide to the subcellular. The number of potential markers probed can be extended even further by staining individual sections with different combinations of antibodies. Thus, this method is flexible enough to allow exploration of varying levels of detail in the sample with many different molecular markers.

Potential application extends beyond preclinical rodent models as human samples have been imaged successfully with LSFM, including developing urogenital organs^44^, the vascular network of the gingiva^45^, and skin biopsies^46^. Importantly in the cardiovascular field, Merz *et al*. successfully imaged a sample of human atrium at single-cell resolution^38^. One consideration during imaging of human tissue is the sample size that can be successfully cleared and fit into the sample chamber. However, the human left anterior descending (LAD) coronary is well within the size that can be successfully imaged. The diameter of a healthy proximal human LAD is 3.7 mm in diameter^47^ and has a wall thickness of 1.0 mm^48^. Hence, we believe that the cardiovascular research potential of this methodology goes beyond preclinical murine models of vascular injury.

In conclusion, we describe a new tool to improve the analysis of preclinical vascular models. LSFM provides increased flexibility in analytical approaches and better precision in injury quantification over traditional histological analysis. Overall, this new workflow based on LSFM and volumetric analysis minimizes bias, it is versatile, more comprehensive than classical histological approaches, and will allow for a deeper understanding of preclinical vasculopathy models. Additionally, we believe this methodology can be applied to the study of other rodent vascular disease models, such as atherosclerosis, hypertension, and aneurysms.

## VI. Acknowledgments

The authors thank Dr. Hussein Kassam for assisting N.E.B. with rat surgeries and Ms. Danielle Berlin for proof reading the manuscript. Microscopy was performed at the UNC Microscopy Services Laboratory. Image analysis was performed at the UNC Neuroscience Center Microscopy Core Facility.

## VII. Author Contributions

E.S.M.B. conceived the idea and directed the work. P.A., E.S.M.B., and N.E.B. established the processing and imaging protocol. N.E.B. drafted the initial manuscript and performed the rat surgeries, sample acquisition, and image analysis. J.L. and F.J.M. performed the mouse surgeries, sample acquisition, and analysis. S.M. performed all histology for the rat arteries. All authors contributed to writing and editing the manuscript.

---

**Video 1.** Injured rat artery slices in x-z and x-y planes followed by volume reconstruction (green = autofluorescence, white = CD31). To see video vitis: https://youtu.be/32oVvm0l3zg

**Video 2.** Injured rat artery slice view in x-y plane followed by volume reconstruction and surface generation of the hyperplastic region. Surface coloring matches that used in Figure 2 (green = autofluorescence, white = CD31, yellow = lumen, blue = internal elastic lamina, pink = external elastic lamina). To see video visit: https://youtu.be/LoAIApmV22o

**Video 3.** Healthy rat artery slices in x-z and x-y planes followed by volume reconstruction (green = autofluorescence, white = CD31). To see video visit: https://youtu.be/fPvTRZzmUy4

