## Supplementary Material (Tables and Figures) for "Light Sheet Fluorescence Microscopy as a New Method for Unbiased Three-Dimensional Analysis of Vascular Injury"

[1]Department of Surgery, Division of Vascular Surgery, University of North Carolina at Chapel Hill, NC 27599.
[2]Center for Nanotechnology in Drug Delivery, University of North Carolina at Chapel Hill, NC 27599.
[3]Curriculum in Toxicology & Environmental Medicine, University of North Carolina at Chapel Hill, NC 27599.
[4]McAllister Heart Institute, University of North Carolina at Chapel Hill, NC 27599
[5]Department of Medicine, Duke University, Durham, NC 27708.
[6]Microscopy Services Laboratory, Department of Pathology and Laboratory Medicine. University of North Carolina at Chapel Hill, NC 27599.
[7]Department of Medicine, Veterans Administration Medical Center, Durham NC 27705.
[8]Department of Cell Biology & Physiology. University of North Carolina at Chapel Hill, NC 27599.

*Corresponding Author:
Edward S. M. Bahnson


### Supplementary Figure 1

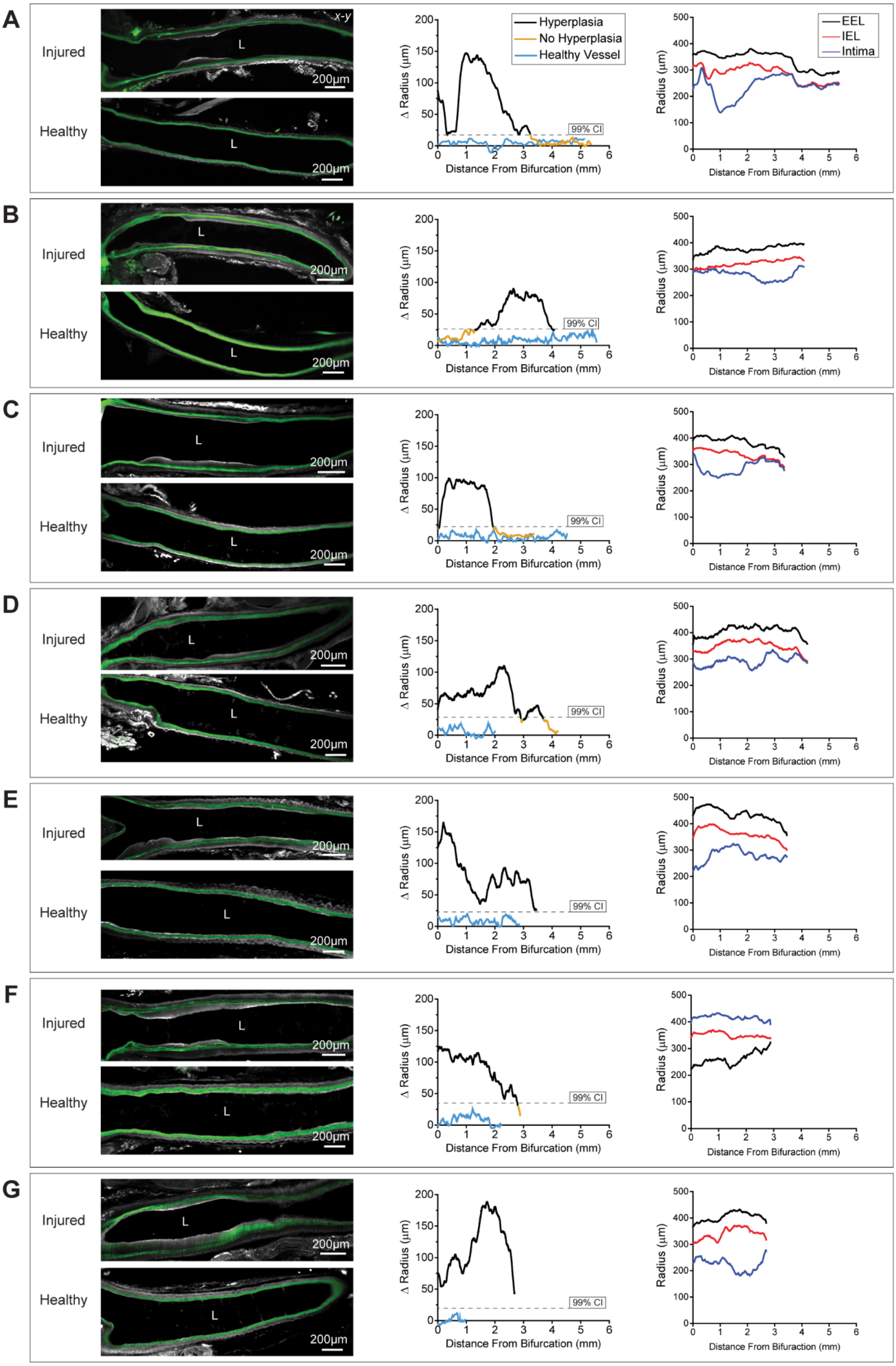

**Supplementary Figure 1.** Representative image and radius analysis of all rat carotid arteries. **(A-G)** Optical slice through the artery center along the x-y axis per injured and healthy rat artery (n = 7, L= lumen, green = autofluorescence, white = CD31). Change (Δ) in radius between the internal elastic lamina and intima radius per artery with 99% confidence interval (CI) hyperplasia threshold identified from the healthy vessel. Last plot depicts the radius from the center of artery to the arterial intima, and internal (IEL) or external (EEL) elastic lamina along the injured artery.

**Supplementary Figure 2**

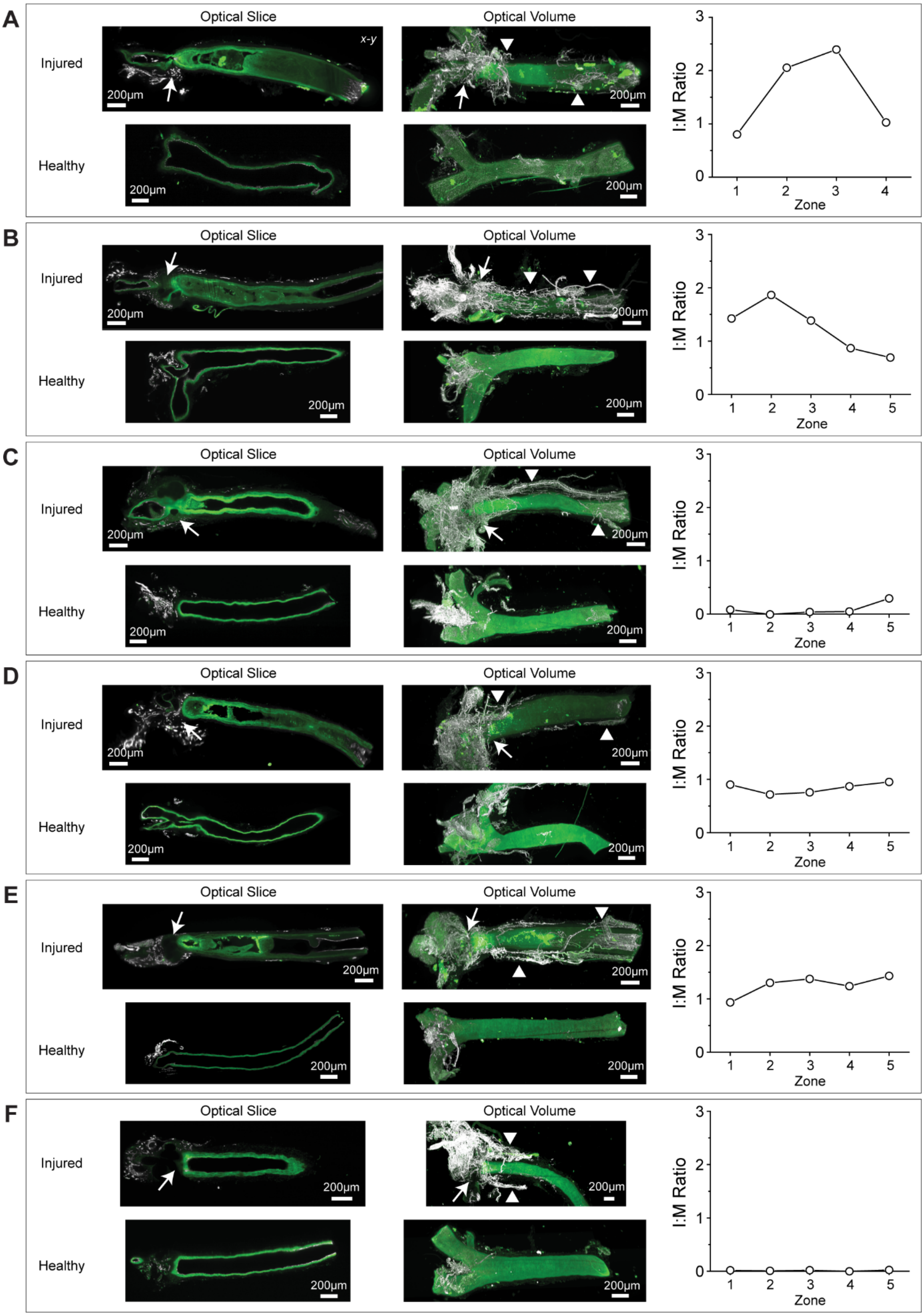

**Supplementary Figure 2.** Representative images of mouse carotid arteries. **(A-F)** Optical slice through the artery center along the x-y axis and volume reconstruction showing periadventitial neovascular plexus following injury (n = 6, arrow = ligation site, arrowhead = regions of neovascular growth, green = autofluorescence, white = CD31). Graph of I:M ratio per injured artery is included. Each zone designates a 500 µm-long segment, with zone 1 starting at the suture site. Average I:M ratio across all zones was used for optical volume value in Figure 5 B and Supplemental Figure 4.

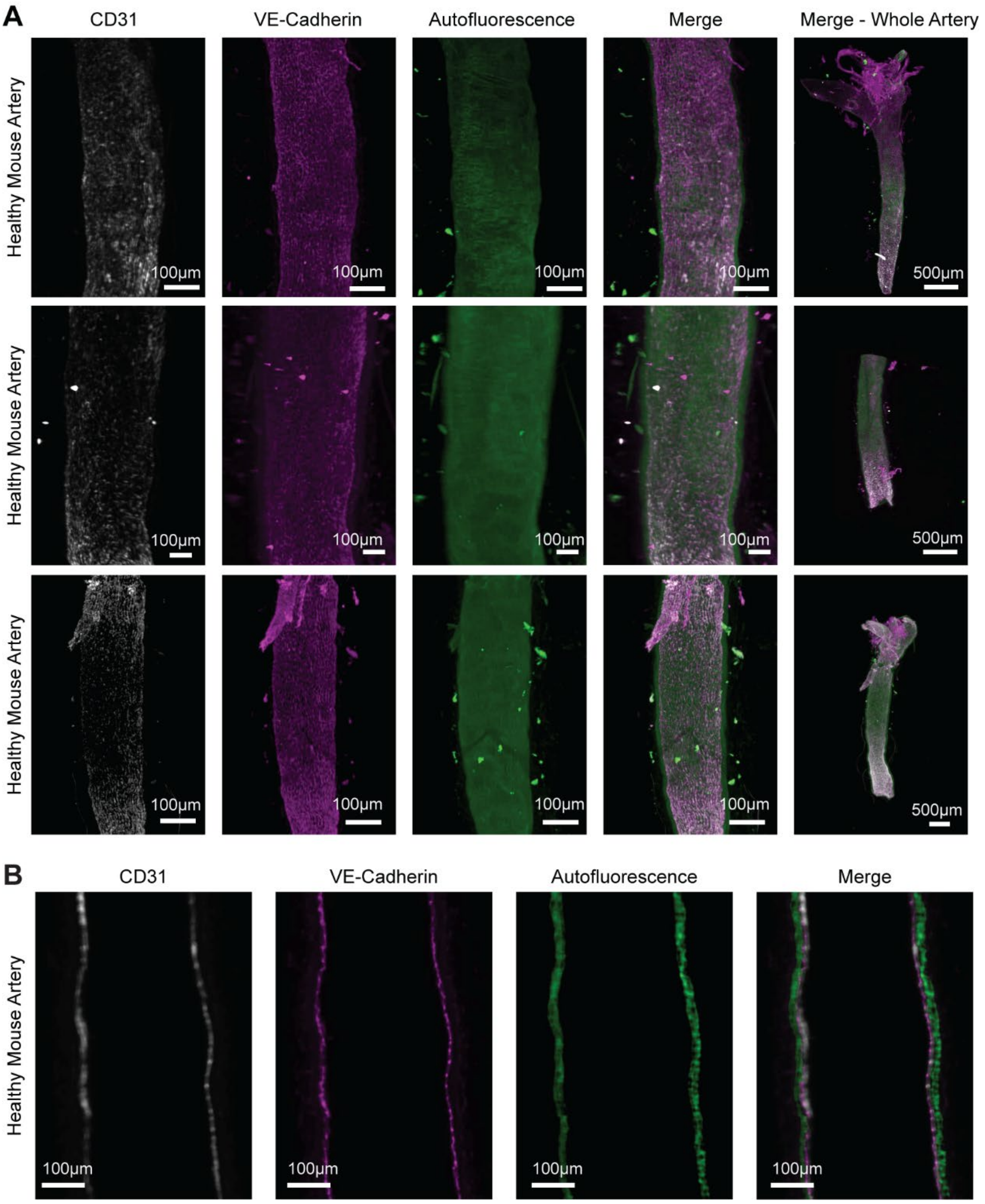


**Supplementary Figure 3.** Multiplex staining of healthy mouse carotid arteries for CD31 and VE-cadherin (n = 3, white = CD31, magenta = VE-cadherin, green = autofluorescence). **A.** 3D rendering of 3 different mouse arteries showing the intact vascular endothelium. **B.** x-y plane longitudinal optical slices showing the endothelial layer of an intact intima, and the elastic lamina of the tunica media

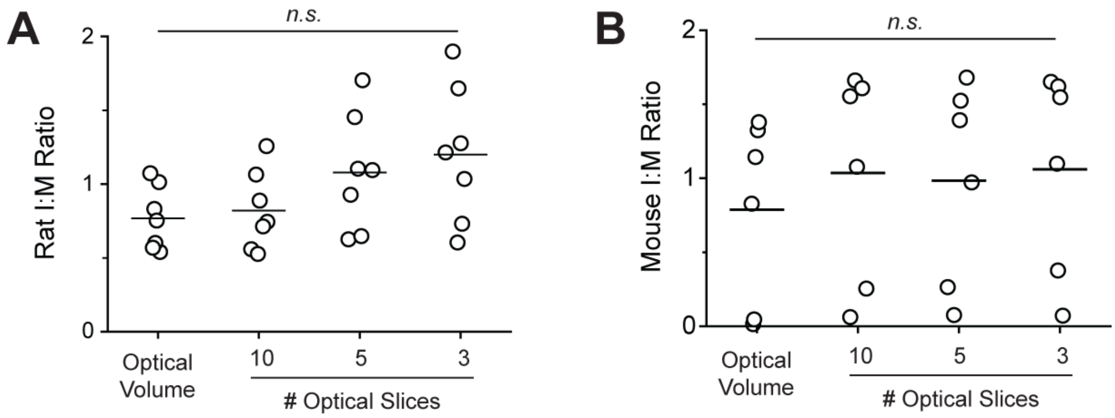


**Supplementary Figure 4.** Performance comparison of optical 3-D volume to optical 2-D slice analysis for obtaining intima-to-media (I:M) ratio. **(A)** I:M ratio obtained per rat using optical volume compared to ImageJ 2-D areal analysis. 3-10 equidistant optical slices in the x-z axis used for 2-D analysis. (n = 7, n.s. = non-significant, one-way ANOVA with Tukey's correction). **(B)** Mouse I:M ratio using optical volume compared to 2-D analysis of 3-10 equidistant optical slices in the x-z axis (n = 6).

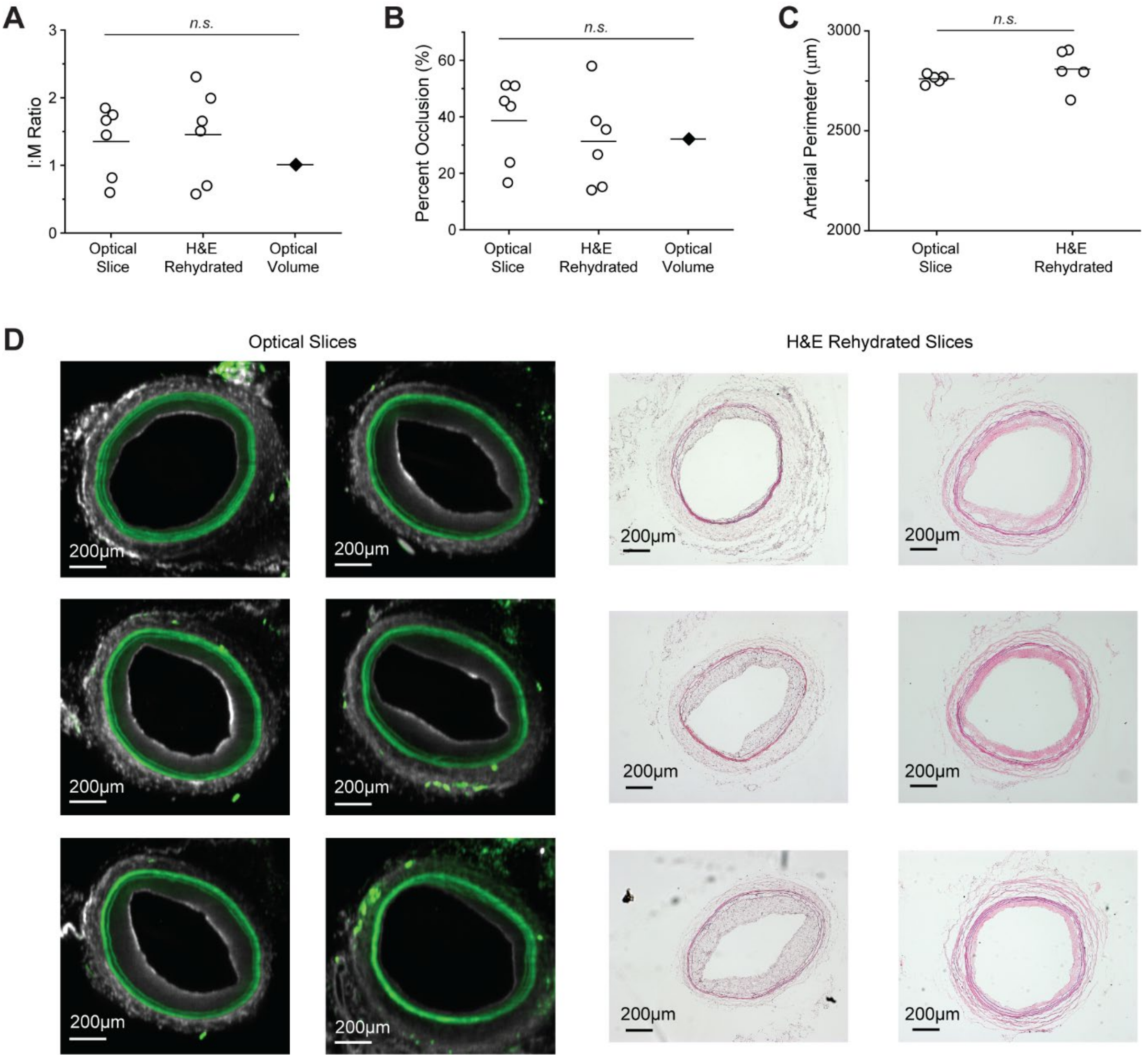


**Supplementary Figure 5.** Performance comparison of optical 3-D volume to optical slice or rehydrated H&E stained 2-D slice analysis for one rat sample. **(A)** Obtained I:M ratio with each point representing a single optical or histological slice (n = 6 slices, n.s. = non-significant, CV% 38% optical slice and 48% histology). **(B)** Obtained percent vessel occlusion values (n = 6 slices, n.s. = non-significant, CV% 38% optical slice and 52% histology). **(C)** Comparison between 2-D histology and optical slice arterial perimeter, as based on the external elastic lamina (n = 5 slices n.s. = non-significant, CV% 1% optical slice and 4% histology). **(D)** Each optical slice or H&E slice used for analysis in (A-C).

**Supplementary Table 1**. Coefficient of variation (CV%) comparing intima:media ratio obtained by 3-D volume to that obtained by 2-D areal analysis.

**CV% for Volume or Designated Number of Slices or Sections**

| Model | Volumetric Analysis | 10 Optical Slices | 5 Optical Slices | 3 Optical Slices | H&E (6-10 Sections) |
|---|---|---|---|---|---|
| Rat Carotid Balloon Injury | 28% | 32% | 37% | 39% | 41% |
| Mouse Carotid Ligation | 79% | 69% | 69% | 65% | 127% |

**Supplementary Table 2**. Results for rat arterial remodeling correlation analysis.

**Neointimal Thickness to Vessel Radius Correlation**

| Lamina | Partial Correlation Coefficient ($\rho_{TR}$) Estimate | Standard Error | 95% CI Lower Bound | 95% CI Upper Bound | P-value |
|---|---|---|---|---|---|
| Internal Elastic Lamina | 0.58 | 0.11 | 0.35 | 0.80 | 4.89E-7 |
| External Elastic Lamina | 0.56 | 0.11 | 0.33 | 0.78 | 1.01E-6 |

### Supplementary Methods

### SAS Code

```
/*Fit Hamlett (2004) mixed model*/
proc mixed data=lsfmdata_long method=ml covtest asycov asycorr;
  class id vtype distance;
  model response = vtype / solution ddfm=kenwardroger;
  random vtype / type=un subject=id v vcorr;
  repeated vtype / type=un subject=distance(id);
  ods output VCorr=VCorr ConvergenceStatus=CS asycorr=asycorr asycov=asycov
CovParms=CovParms;
run;
/*Use delta method to find SE(rho), calculate CI/p-value based on
normality*/
proc iml;
  use Covparms;
  read all var {Estimate} into Cov_estimate;
  close Covparms;
  use Asycov;
  read all var {CovP1 CovP2 CovP3 CovP4 CovP5 CovP6} into asycov;
  close Asycov;
  a = Cov_estimate[1];
  b = Cov_estimate[2];
  c = Cov_estimate[3];
  g = Cov_estimate[4];
  h = Cov_estimate[5];
  i = Cov_estimate[6];
  rhohat = (b+h)/sqrt((a+g)*(c+i));
  df_da = -0.5*((b+h)*(c+i))/sqrt(((a+g)*(c+i))**3);
  df_db = 1/sqrt((a+g)*(c+i));
  df_dc = -0.5*(b+h)*(a+g)/sqrt(((a+g)*(c+i))**3);
  df_dg = -0.5*(b+h)*(c+i)/sqrt(((a+g)*(c+i))**3);
  df_dh = 1/sqrt((a+g)*(c+i));
  df_di = -0.5*(b+h)*(a+g)/sqrt(((a+g)*(c+i))**3);
  create partialderiv var {df_da df_db df_dc df_dg df_dh df_di};
  append;
  close partialderiv;
  use partialderiv;
  read all var {df_da df_db df_dc df_dg df_dh df_di} into partialderiv;
  close partialderiv;
  Sigma = asycov;
  rho_var = partialderiv*Sigma*t(partialderiv);
  rho_SE = sqrt(rho_var);
  l95=rhohat-1.96*rho_SE;
  u95=rhohat+1.96*rho_SE;
  p = (1-cdf("normal",abs(rhohat),0,rho_SE))*2;
  NormalApproxOutput = j(1,5,.);
  NormalApproxOutput[,1]=rhohat;
  NormalApproxOutput[,2]=rho_SE;
  NormalApproxOutput[,3]=l95;
  NormalApproxOutput[,4]=u95;
  NormalApproxOutput[,5]=p;
  Outtitle={'Estimate' 'Std Error' '95% CI Lower Bound' '95% CI Upper
Bound' 'Pvalue'};
  Varnames=t("intimal thickness and vessel radius");
  PRINT NormalApproxOutput[colname=Outtitle rowname=Varnames];
quit;
```
